# Distinct Roles for the Thioredoxin and Glutathione Antioxidant Systems in Nrf2-Mediated Lung Tumor Initiation and Progression

**DOI:** 10.1101/2024.08.20.608800

**Authors:** Amanda M. Sherwood, Janine M. DeBlasi, Samantha Caldwell, Gina M. DeNicola

## Abstract

Redox regulators are emerging as critical mediators of lung tumorigenesis. NRF2 and its negative regulator KEAP1 are commonly mutated in human lung cancers, leading to NRF2 accumulation and constitutive expression of NRF2 target genes, many of which are at the interface between antioxidant function and anabolic processes that support cellular proliferation. Nrf2 activation promotes lung tumor initiation and early progression in murine models of lung cancer, but which Nrf2 targets mediate these phenotypes is unknown. Nrf2 regulates two parallel antioxidant systems mediated by thioredoxin reductase 1 (TXNRD1) and glutathione reductase (GSR), which promote the reduction of protein antioxidant thioredoxin (TXN) and tripeptide antioxidant glutathione (GSH), respectively. We deleted TXNRD1 and GSR alone, or in combination, in lung tumors harboring mutations in Kras^G12D^ and Nrf2^D29H^. We found that tumor initiation was promoted by expression of GSR, but not TXNRD1, regardless of Nrf2 status. In contrast, Nrf2^D29H^ tumors, but not Nrf2^WT^, were dependent on TXNRD1 for tumor progression, while GSR was dispensable. Simultaneous deletion of GSR and TXNRD1 reduced initiation and progression independent of Nrf2 status, but surprisingly did not completely abrogate tumor formation. Thus, the thioredoxin and glutathione antioxidant systems play unique roles in tumor initiation and progression.

## Introduction

The transcription factor NRF2 (nuclear factor-erythroid 2 p45-related factor 2) is a stress-responsive transcription factor that regulates the cellular antioxidant response^1,2^ and other facets of cellular metabolism^3^. KEAP1 (Kelch-like ECH-associated protein 1) is a negative regulator of NRF2 and under non-stressed conditions, facilitates the ubiquitin-mediated degradation of NRF2^4^. Reactive oxygen species and other electrophiles modify reactive cysteine residues on KEAP1, resulting in impaired NRF2 degradation and translocation of NRF2 into the nucleus^5,6^, thereby promoting the expression of cytoprotective genes^2,7–14^.

The two main antioxidant systems regulated by NRF2 are the glutathione (GSH)^2,7,15–19^ and the thioredoxin (TXN) systems^7,19–21^. The GSH system consists of the tripeptide antioxidant glutathione, which is used by glutathione peroxidases (GPXs)^22,23^ in the defense against lipid peroxides^22,24,25^. The pool of GSH is maintained by both de novo synthesis and by glutathione reductase (GSR)^16^, which regenerates GSH from its oxidized form GSSG. The TXN system is comprised of peroxiredoxins (PRXs)^26^, which are the main defense mechanism against hydrogen peroxides^27,28^. Oxidized PRXs are reduced by the protein antioxidant thioredoxins (TXNs)^26,29^, which are subsequently reduced by thioredoxin reductases (TXNRDs)^9,28,30^. Both systems overlap in function and targets. GSR and TXNRD1 reduce disulfide bonds and convert cystine (Cys2) to cysteine (Cys)^31–34^. The TXN system can also compensate for GSH deficiency, suggesting semi-redundant roles for these systems^32^. Despite their overlap, these redox systems have distinct ROS-scavenging mechanisms^35–37^ and differentially regulate cell signaling^37–40^.

We and others have previously reported on the oncogenic role of Nrf2 in Kras^G12D^-mediated lung tumor initiation and progression^14,41–49^. However, the roles of the Nrf2-regulated antioxidant systems in mediating these phenotypes remains unknown. We focused on TXNRD1 and GSR, the key mediators of these systems. We utilized the Kras^G12D/+^ model of early lung adenocarcinoma^50^, in which tumors expressed either Nrf2^WT^ or a Nrf2^D29H/+^ mutation to promote Nrf2 stabilization^49^. Using this cancer model, we deleted GSR or TXNRD1 alone or in combination to study the intrinsic roles of these systems in tumor initiation and progression.

## Results

### Nrf2 promotes the expression of GSR and TXNRD1 in murine lung tumors

We first determined whether Nrf2 activation increased the protein expression of GSR and TXNRD1 in lung tumors by performing immunohistochemical (IHC) staining for the Nrf2 targets NQO1, GSR and TXNRD1 in Kras^G12D^; Nrf2^+/+^ and Kras^G12D^; Nrf2^D29H^ lung tumors. As previously reported, Nrf2^D29H^ increased the expression of NQO1 in tumors (Figs. 1A-B)^49^ and also increased the expression of GSR and TXNRD1 (Figs. 1C–F). While bronchiolar hyperplasias had higher expression of these proteins compared to alveolar hyperplasias and grade 1 and grade 2 adenomas, Nrf2^D29H^ promoted their expression across most lesion types and expression remained high across tumor grades (Figs. 1B, D, F). These results indicate that Nrf2 promotes GSR and TXNRD1 expression in Kras^G12D/+^ lung tumors *in vivo*, and suggests these proteins may mediate the effects of Nrf2^D29H^ in promoting initiation and progression.

**Figure 1.**
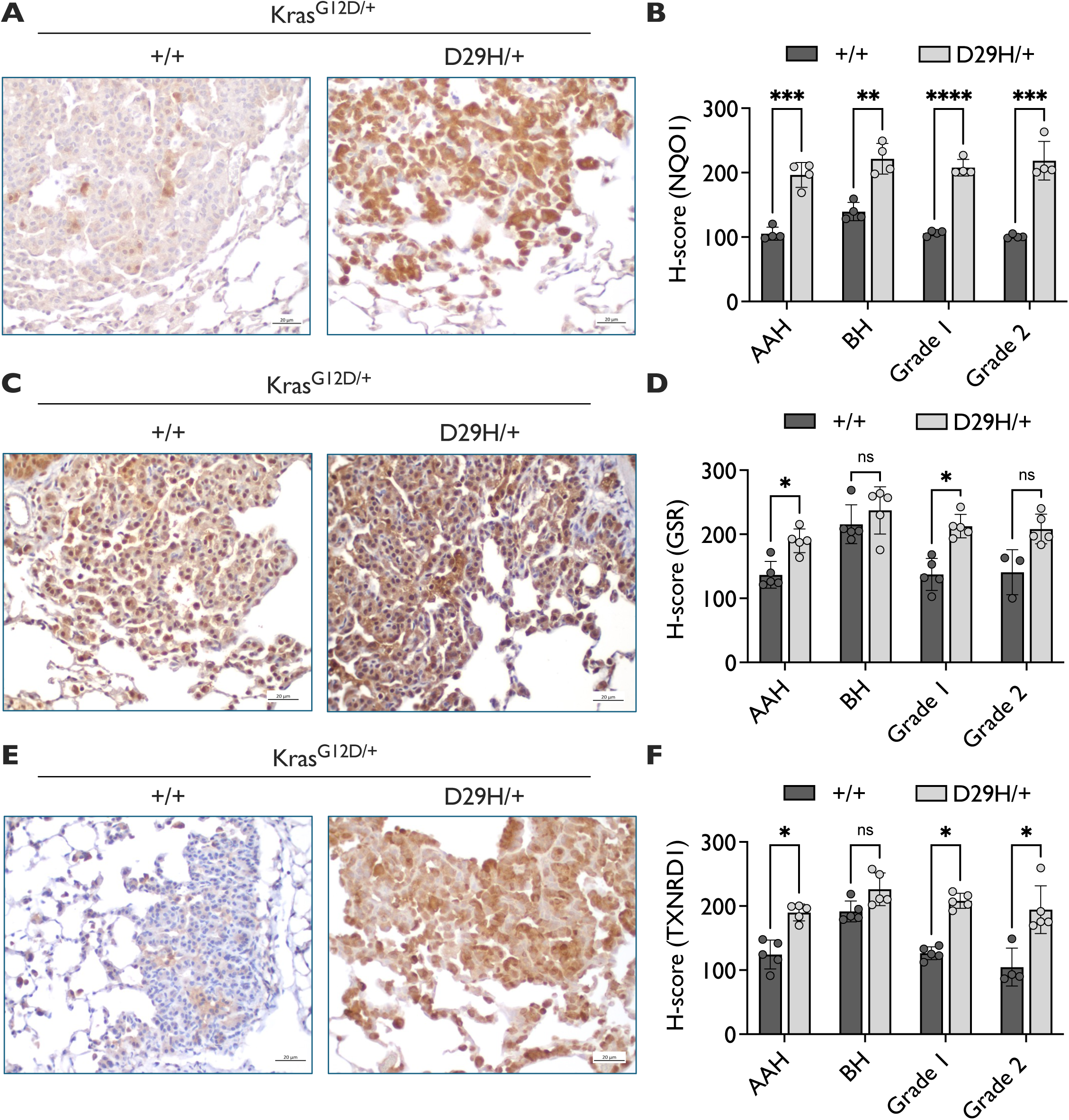
NQO1, GSR and TXNRD1 expression increase upon Nrf2 activation. Representative IHC staining for (**A**) NQO1, (**C**) GSR and (**E**) TXNRD1 in Kras^G12D/+^ and Kras^G12D/+;^ Nrf2^D29H/+^ tumors. Scale bars, 20 μm. Graphs depicting the H-score per grade in Nrf2^+/+^ and Nrf2^D29H/+^ tumors for (**B**) NQO1 (**D**) GSR and (**F**) TXNRD1. Nrf2^+/+^ (*n* = 4); Nrf2^D29H/+^ (*n* = 4) in (B); Nrf2^+/+^ (*n* = 5, except where sufficient tumors were available for grade 2 analysis); Nrf2^D29H/+^ (*n* = 5) in (D) and (F). Since grade 3 tumors are rare in the Kras^G12D/+^ model, the H-score analysis per grade was only conducted up to grade 2 tumors. *p<0.05, **p<0.01, ***p<0.001, ****p<0.0001 (unpaired t-test with Holm–Sidak multiple comparisons test). AAH, alveolar adenomatous hyperplasia; BH, bronchiolar hyperplasia.

### GSR deficiency impairs lung tumor initiation, but not progression, independent of Nrf2 status

Using the Gr1^a1Neu^ allele^51,52^ which is a functional knockout (KO) of GSR, we generated Kras^G12D/+^; Nrf2^+/+^; Gsr^a1Neu/a1Neu^ (GSR KO) and Kras^G12D/+^; Nrf2^D29H/+^; Gsr^a1Neu/a1Neu^ (GSR KO) lung tumors to evaluate the requirement of GSR for tumor initiation and progression. First, we validated the lack of GSR protein by performing immunohistochemical staining for GSR (Fig. 2A-B). To examine the influence of GSR on tumor initiation, we quantified tumor number across the genotypes and found that GSR KO significantly decreased tumor number in both the Kras^G12D/+^; Nrf2^+/+^ and Kras^G12D/+^; Nrf2^D29H/+^ models (Figs. 2C–D). We next examined the impact of GSR on tumor progression by analyzing tumor grade from adenomatous and bronchiolar hyperplasia (AAH and BH, respectively) to tumors from grades 1 (adenoma) to 3 (adenocarcinoma). However, GSR expression did not impact lung tumor grade, and Nrf2^D29H/+^ still promoted an increase in grade 1 tumors as we previously reported (Fig. 2E)^49^. Interestingly, we did see an impairment of grade 3 tumor formation in the Nrf2^D29H/+^ GSR WT and Nrf2^D29H/+^ GSR KO tumors, consistent with our previous study showing that Nrf2 activation blocks progression to advanced grade tumors^49^. We also examined lung tumor burden by grade, which is defined as the percentage of the lung covered by each tumor grade. Interestingly, while GSR did not influence grade 1 tumor burden in the Nrf2^+/+^ model, it did decrease grade 1 tumor burden and size in the Nrf2^D29H/+^ model (Fig. S1A-B). Surprisingly, GSR loss also decreased grade 2 tumor burden and size, regardless of Nrf2 status (Fig. S1A-B). To test if GSR KO resulted in compensatory induction of Nrf2 and TXNRD1, we performed immunohistochemistry to stain for TXNRD1 and NQO1. However, we found that GSR KO did not alter the expression these proteins (Fig. 2F, Fig. S2A-B). Overall, these findings indicate that while GSR promotes the formation of lung tumors, GSR is dispensable for the early progression to low-grade tumors, but may contribute to their growth.

**Figure 2.**
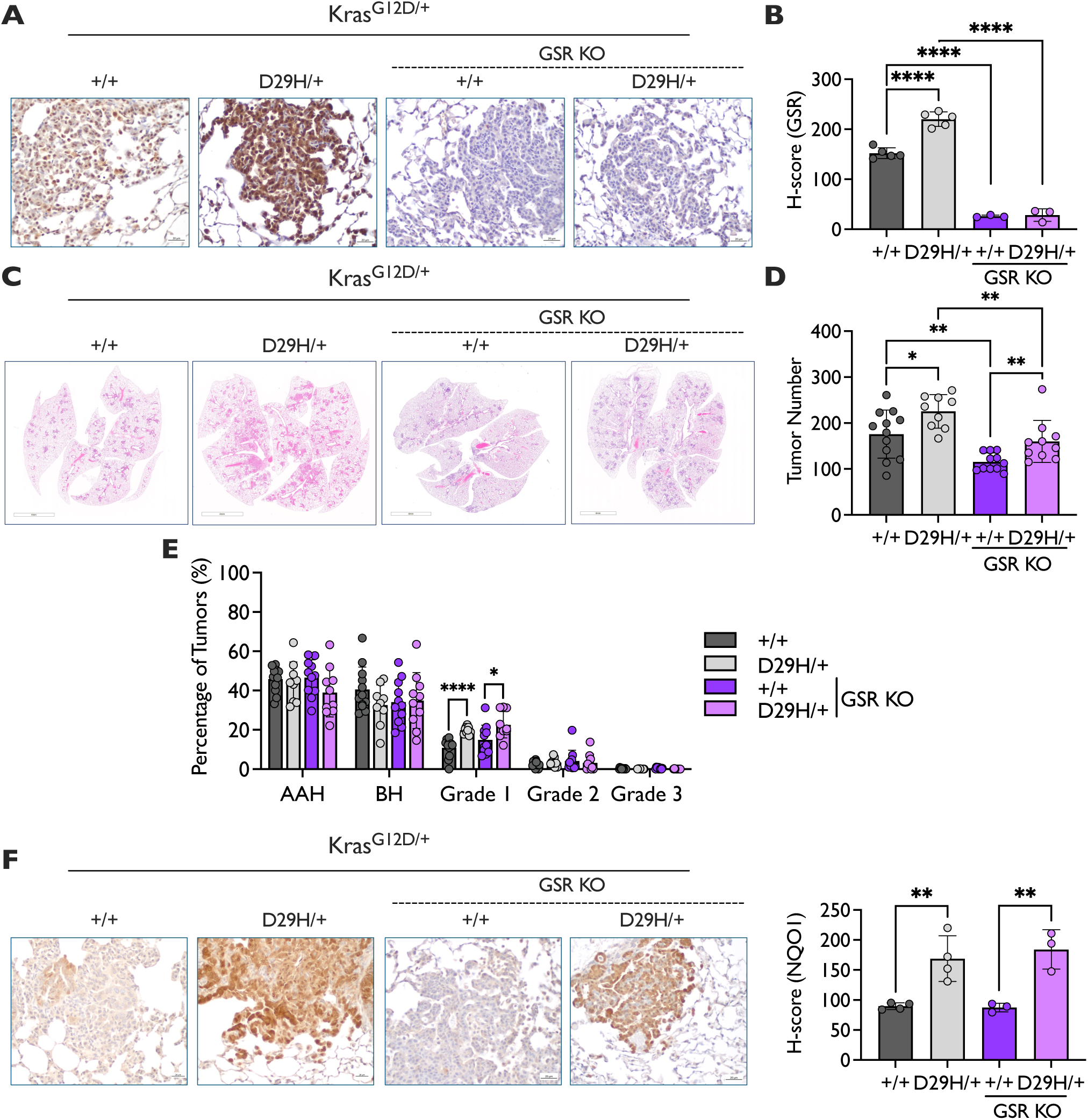
GSR KO impairs Nrf2^+/+^ and Nrf2^D29H/+^ tumor initiation. **(A)** Representative GSR immunohistochemical (IHC) staining of Kras^G12D/+^ and Kras^G12D/+;^ Nrf2^D29H/+^ tumors. Scale bars, 20 μm. **(B)** H-scores for GSR IHC staining. GSR WT (*n* = 5), GSR KO (*n* = 3). **(C)** Representative whole-lung hematoxylin and eosin–stained section. Scale bars, 4,000 μm. **(D)** Tumor number per mouse of GSR KO mice (*n* = 11 for Nrf2^+/+^; *n* = 10 for Nrf2^D29H/+^) compared to GSR WT mice (*n* = 12 for Nrf2^+/+^; *n* = 9 for Nrf2^D29H/+^). *p<0.05, **p<0.01. Unpaired t-test. **(E)** Distribution of tumor grades across GSR WT (*n* = 12 for Nrf2^+/+^; *n* = 9 for Nrf2^D29H/+^) and GSR KO (*n* = 11 for Nrf2^+/+^; *n* = 10 and Nrf2^D29H/+^) models. *p<0.05, ****p<0.0001 (unpaired t-test with Holm–Sidak multiple comparisons test). **(F)** Representative IHC staining of NQO1 in Kras^G12D/+^ and Kras^G12D/+;^ Nrf2^D29H/+^ mice. Scale bars, 20 μm. H-scores for NQO1 IHC staining. GSR WT (*n* = 4), GSR KO (*n* = 3).

### TXNRD1 deficiency impairs lung Nrf2^D29H^-dependent tumor progression, but not initiation

We next focused on TXNRD1. Using the Txnrd1^flox^ allele^53^, we generated Kras^G12D/+^; Nrf2^+/+^; Txnrd1^Λ/Λ^ (TXNRD1 KO) and Kras^G12D/+^; Nrf2^D29H/+^; Txnrd1^Λ/Λ^ (TXNRD1 KO) lung tumors to evaluate the requirement of TXNRD1 for tumor initiation and progression. We performed immunohistochemical staining of TXNRD1 protein to confirm that TXNRD1 was knocked out (Fig.3A-B). Both TXNRD1 KO models were largely negative for TXNRD1, although there was still some patchy expression largely localized to macrophages and some bronchiolar cells. Unlike what was observed upon GSR loss, we found that knocking out TXNRD1 had no impact on tumor number in the Kras^G12D/+^, Nrf2^+/+^ mice or Kras^G12D/+^, Nrf2^D29H/+^ mice (Figs. 3C–D). We then examined the role of TXNRD1 on tumor progression by analyzing tumor grades. TXNRD1 KO abolished the Nrf2^D29H^-mediated increase in grade 1 adenomas (Fig. 3E), but had no effect on Nrf2^+/+^ tumor progression to grade 1 adenomas, although there was a small decrease in the percentage of grade 2 tumors (Fig. 3E), although these tumors are rare in this model. Surprisingly, there was a significant increase in the burden of Nrf2^+/+^ grade 1 tumors upon TXNRD1 KO, which was accompanied by an increase in grade 1 tumor size (Fig. S1C–D). Because we noticed TXNRD1 KO tumors appeared to be heavily infiltrated by macrophages upon macroscopic examination of the H&E images, we performed immunohistochemical staining for the macrophage marker F4/80. We found that there was a significant increase in F4/80 staining in TXNRD1 KO tumors compared to all other genotypes, with as much as 40% of the tumor content being macrophages (Fig. S3). Thus, the increase in grade 1 burden in Nrf2^+/+^ TXNRD1 KO tumors may be accounted for by the enhanced macrophage infiltration. Like GSR loss, TXNRD1 loss did not affect the expression of the Nrf2 target proteins NQO1 (Fig. 3F) and GSR (Fig. S2C-D), suggesting that inactivating the TXN system does not activate Nrf2 or induce compensatory GSR upregulation. Overall, these findings indicate that although TXNRD1 is dispensable for tumor initiation, TXNRD1 mediates the tumor progression effects of Nrf2^D29H/+^. Furthermore, the TXN system may play an immunosuppressive role and disruption of this system leads to increased macrophage infiltration.

**Figure 3.**
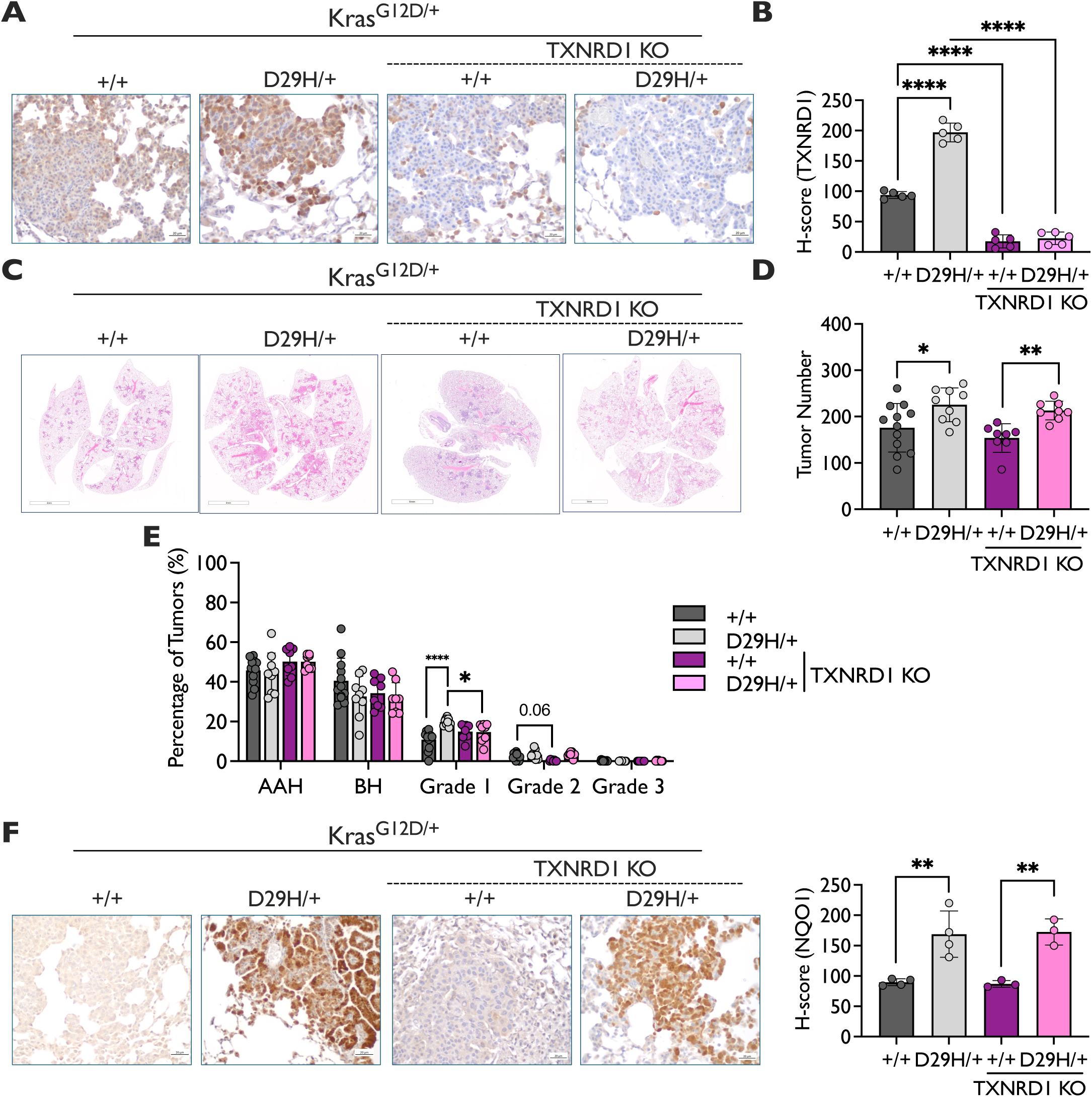
TXNRD1 KO impairs NRF2^D29H/+^ tumor progression. **(A)** Representative TXNRD1 immunohistochemical (IHC) staining of Kras^G12D/+^ and Kras^G12D/+;^ Nrf2^D29H/+^ tumors. Scale bars, 20 μm. **(B)** H-scores for TXNRD1 IHC staining. *n* = 5 mice per genotype. **(C)** Representative whole-lung hematoxylin and eosin–stained section. Scale bars, 4,000 μm. **(D)** Tumor number per mouse of TXNRD1 KO mice (*n* = 8 for Nrf2^+/+^; *n* = 8 for Nrf2^D29H/+^) compared to TXNRD1 WT mice (*n* = 12 for Nrf2^+/+^; *n* = 9 for Nrf2^D29H/+^). N.B. TXNRD1 WT mice are the same as GSR WT mice from Fig. 2D. *p<0.05, **p<0.01. Unpaired t-test. (**E**) Distribution of tumor grades across TXNRD1 WT (*n* = 12 for Nrf2^+/+^; *n* = 9 for Nrf2^D29H/+^). and TXNRD1 KO mice (*n* = 8 for Nrf2^+/+^; *n* = 8 for Nrf2^D29H/+^) models. N.B. TXNRD1 WT mice are the same as GSR WT mice from Fig. 2E. *p<0.05, **p<0.01, ****p<0.0001 (unpaired t-test with Holm–Sidak multiple comparisons test). **(F)** Representative NQO1 IHC staining of Kras^G12D/+^ and Kras^G12D/+;^ Nrf2^D29H/+^ mice. Scale bars, 20 μm. H-scores for NQO1 IHC staining. TXNRD1 WT (*n* = 4), TXNRD1 KO (*n* = 3).

### Simultaneous ablation of GSR and TXNRD1 impairs initiation and progression

The differential effects on tumor initiation and progression following GSR and TXNRD1 ablation in Kras^G12D/+^ tumors suggests non-overlapping functions; however, in both models, tumors were still able to form. GSR and TXNRD1 are the only two disulfide reductases in the cytoplasm and concomitant deletion of both would be expected to prevent the reduction of glutathione, protein disulfides, and cystine, and thereby induce lethal oxidative stress. To address the consequence of full disulfide reductase loss in tumors, we generated Kras^G12D/+^; Nrf2^+/+^ and Kras^G12D/+^; Nrf2^D29H/+^ TXNRD1/GSR double KO (DKO) tumors. We performed immunohistochemical staining of TXNRD1 protein to confirm that TXNRD1 was deleted in the DKO model (Figs. 4A–C). Unlike the TXNRD1 KO model, which was largely negative for TXNRD1, the DKO model had staining for TXNRD1, although it was less than the TXNRD1 WT model, suggesting some tumors had fully recombined the TXNRD1^flox^ allele (Fig. 4B). We analyzed what fraction of tumors were TXNRD1 positive and found that roughly 40% of tumors in both the Kras^G12D/+^; Nrf2^+/+^ and Kras^G12D/+^; Nrf2^D29H/+^ models had retained TXNRD1 expression (Fig. 4C). Thus, lung tumors lacking both GSR and TXNRD1 could form, although escape from TXNRD1 recombination was frequent, suggesting loss of both disulfide reductases was disadvantageous.

**Figure 4.**
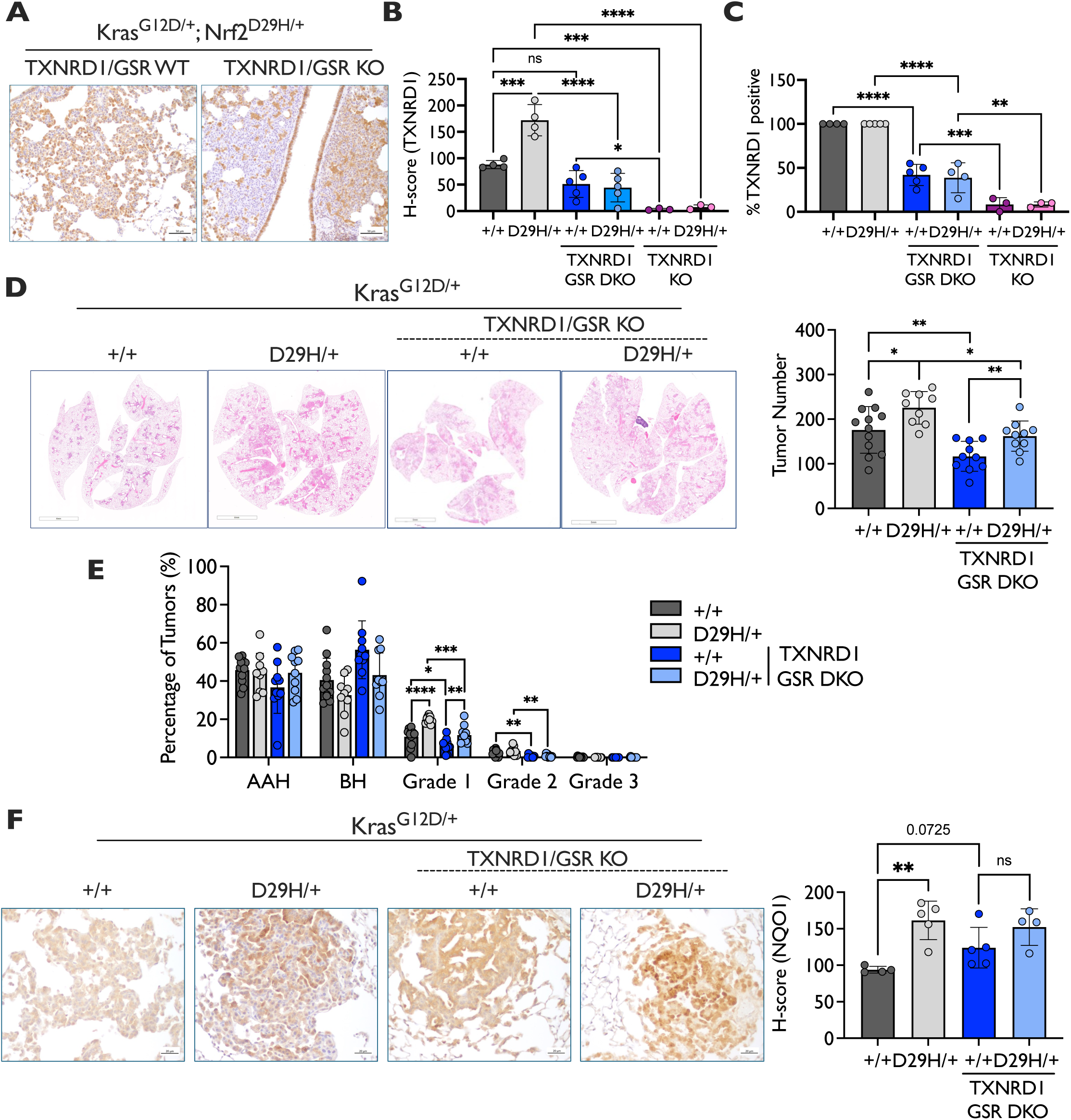
Dual deletion of GSR and TXNRD1 impairs Nrf2^+/+^ and Nrf2^D29H/+^ tumor formation and progression. (**A**) Representative TXNRD1 IHC staining of Kras^G12D/+;^ Nrf2^D29H/+^ TXNRD1/GSR WT and DKO tumors depicting positive and negative tumors. Scale bars, 50 μm. (**B**) H-score of TXNRD1 expression and (**C**) % TXNRD1 positive tumors quantification of Kras^G12D/+;^ Nrf2^+/+^ and Kras^G12D/+;^ Nrf2^D29H/+^ GSR/TXNRD1 WT (*n* =4), TXNRD1 KO (*n* =3) and GSR/TXNRD1 DKO mice (*n* =5). *p<0.05, **p<0.01, ***p<0.001, ****p<0.0001. Unpaired t-test. (**D**) Representative whole-lung hematoxylin and eosin–stained sections. Scale bars, 4,000 μm. Tumor number per mouse of TXNRD1/GSR DKO mice (*n* = 10 for Nrf2^+/+^ and Nrf2^D29H/+^) compared to TXNRD1/GSR WT mice (*n* = 12 for Nrf2^+/+^; *n* = 9 for Nrf2^D29H/+^). N.B. TXNRD1/GSR WT mice are the same as GSR WT mice from Fig. 2D. *p<0.05, **p<0.01. Unpaired t-test. (**E**) Distribution of tumor grades across TXNRD1/GSR WT mice (*n* = 12 for Nrf2^+/+^; *n* = 9 for Nrf2^D29H/+^) and TXNRD1/GSR DKO mice (*n* = 10 for Nrf2^+/+^ and Nrf2^D29H/+^) models. N.B. TXNRD1/GSR WT mice are the same as GSR WT mice from Fig. 2E. *p<0.05, **p<0.01, ***p<0.001, ****p<0.0001 (unpaired t-test with Holm–Sidak multiple comparisons test). **(F)** Representative NQO1 immunohistochemical (IHC) staining of Kras^G12D/+^ and Kras^G12D/+;^ Nrf2^D29H/+^ mice. Scale bars, 20 μm. H-scores for NQO1 IHC staining. **p<0.01. Unpaired t-test. TXNRD1/GSR WT (*n* = 4 for Nrf2+/+ and *n* = 5 for Nrf2^D29H/+^); TXNRD1/GSR KO (*n* = 5 for Nrf2+/+ and *n* = 4 for Nrf2^D29H/+^).

Like what we observed in the GSR KO model, we found that TXNRD1/GSR DKO significantly reduced tumor formation in both the Kras^G12D/+^, Nrf2^+/+^ mice and Kras^G12D/+^, Nrf2^D29H/+^ models (Fig. 4D). We next examined the effect of TXNRD1/GSR DKO on tumor progression. Unlike TXNRD1 KO, which selectively influenced Nrf2^D29H/+^ progression, TXNRD1/GSR DKO decreased the percentage of grade 1 tumors in both the Nrf2^+/+^ and Nrf2^D29H/+^ models (Fig. 4E). This was accompanied by a decrease in grade 1 tumor burden and tumor size in TXNRD1/GSR DKO mice compared to TXNRD1/GSR WT mice (Fig. S1E–F). We also observed a decrease in grade 2 percentage, burden, and size in the Nrf2^+/+^ model, and an impairment in grade 3 tumor formation in both the Nrf2^+/+^ and Nrf2^D29H/+^ models following GSR/TXNRD1 KO (Figs. 4E, S1E– F). We also performed immunohistochemical staining for the Nrf2 target NQO1 to see if impairing both the GSH and TXN systems resulted in increased Nrf2 activity. Interestingly, there was a modest increase in NQO1 activation in the Nrf2^+/+^ TXNRD1/GSR DKO tumors (Fig. 4F), suggesting loss of both enzymes induces oxidative stress. Overall, these data suggest that the GSH and TXN systems have different, although complementary, roles in initiation and progression and that inhibition of both systems impairs tumor formation and progression.

## Discussion

Our finding that the GSH system was more essential for tumor initiation while the TXN system was more essential for tumor progression is consistent with prior work in breast cancer, which found that inhibition of GSH synthesis impaired tumor initiation, but once formed tumors became reliant on the TXN system for growth^54^. However, our study differs from this study in that they inhibited GSH synthesis, whereas we targeted GSH regeneration. Given that our tumors were still capable of GSH synthesis, it is surprising that inhibition of GSH recycling impaired tumor initiation significantly. Because Nrf2^D29H^ still promoted tumor initiation in the absence of GSR, it is likely that this is due to expression of GCLC and GCLM, Nrf2 targets that promote GSH synthesis^17,19^. Future work combining GSR KO with inhibition of GSH synthesis will reveal whether inhibition of both GSH synthesis and recycling abolishes the Nrf2-mediated tumor initiation phenotype.

We found that not only was TXNRD1 essential for Nrf2^D29H^-mediated tumor progression, but TXNRD1 KO tumors also exhibited an increase in macrophage infiltration. One potential explanation for this is ER stress, which induces cytokine production that may promote macrophage recruitment^55^: TXNRD1 is required for protein folding in the ER^56^, and ER stress has been linked to glycosylation of cytokines and immune cell recruitment^57^. Secreted TXN also displays cytokine-like properties^58,59^, serving as a chemoattractant for various immune cell types^60^. Additional work is needed to understand why macrophage infiltration is increased in TXNRD1 KO tumors, whether these macrophages are protumorigenic or antitumorigenic, and why these macrophages are not present in TXNRD1/GSR DKO tumors.

Tumors lacking both TXNRD1 and GSR could unexpectedly form but it is unclear how this is possible. TXNRD1/GSR null hepatocytes maintain redox homeostasis by shuttling dietary methionine into the transsulfuration pathway to produce Cys for GSH synthesis^61^. However, we found that both healthy mouse lungs and LUAD tumors lack transsulfuration capacity for cysteine synthesis^62^ and thus should not be able to sustain their redox state from methionine. Additional research is needed to identify other antioxidant systems or cysteine acquisition mechanisms compensating for the loss of GSR and TXNRD1.

Lastly, although we and others have shown that Nrf2 promotes lung tumor initiation and early progression, we previously reported that Nrf2 activation blocks progression to advanced grade adenocarcinomas^49^, which is also evident in this study in which Nrf2^D29H^ mice lack grade 3 tumors (Fig. S1A). Chronic Nrf2 stabilization has been associated with various metabolic liabilities, including reductive stress^63^, glutamate deprivation^14,64^, and CDO1-dependent NADPH deletion^11^. Future work in GEMMs of advanced LUAD progression such as the LSL-Kras^G12D/+^; p53^flox/flox^ model will reveal whether the glutathione and thioredoxin systems play a role in Nrf2-mediated adenocarcinoma suppression. Moreover, they will be critical in clarifying the roles of the GSH and TXN systems in advanced LUAD independently of Nrf2.

## Materials and Methods

### Mice

The housing and breeding of mice in this study adhered to the ethical guidelines and approval of the Institutional Animal Care and Use Committee (protocols #: IS00007922R). Alleles in this study were described previously, including the *LSL-Nfe2l2^D29H^* (MGI: 7327101)^49^; *Txnrd1^flox^* (RRID: IMSR_JAX:028283)^53^, *ROSA^mT/mG^* (RRID:IMSR_JAX:007576)^65^, *LSL-Kras^G12D/+^* (RRID:IMSR_JAX:008179)^50^ and *Gr1^a1Neu^* (MGI: 1857772)^51,52^ alleles. All mice were kept on a mixed C57BL/6 genetic background. Mice were infected intranasally with 2.25 × 10^7^ PFU adenoviral-Cre to induce lung tumors under anesthesia as previously described^49^.

### Immunohistochemistry

Paraffin-embedded tumor-bearing mouse lungs were sectioned, deparaffinized and rehydrated, followed by heat-mediated antigen retrieval with boiling 10 mmol/L citrate buffer (pH 6.0). Antibodies used for IHC are as follows: GSR (Santa Cruz Biotechnology, Cat # sc-133245, 1:100), TXNRD1 (Cell Signaling Technologies, Cat #15140S), NQO1 (Sigma-Aldrich, Cat #HPA007308, 1:250), and F4/80 (Cell Signaling Technologies, 1:250). Sections were incubated at 4°C in primary antibody overnight. The ImmPRESS HRP (horseradish peroxidase) goat anti-rabbit kit (Vector Laboratories, RRID:AB_2631198) was used as directed by the manufacturer’s instructions. For the mouse antibody GSR, the M.O.M (Mouse-on-Mouse) ImmPRESS IgG HRP polymer kit (Vector Laboratories, RRID:AB_2336832) was used to reduce endogenous mouse Ig staining. DAB peroxidase (HRP) substrate (Vector Laboratories, SK-4105) was used, followed by counterstaining with hematoxylin (Vector Laboratories, H-3404). Slide scanning at 20× was performed with an Aperio imager and the H-score of at least five representative regions/mouse was analyzed with QuPath software (Bankhead et al, Qupath Sci Rep 2017). The Axio Lab A.1 microscope was used to take representative images at x 40 (Carl Zeiss Microimaging Inc.)

### Tumor grading analysis and histology

Lung tumors were manually graded using a previously published protocol^50^. Tumor number was determined by normalizing the total tumor number per mouse H&E section to the lung area. The distribution of tumor grades among genotypes was determined by dividing the number of tumors per grade by the number of tumors per mouse. Tumor burden per grade was obtained by dividing the area of the lung occupied by tumors of a specific grade by the total lung area.

### Quantification and statistical analysis

GraphPad Prism9 software was used to perform unpaired t-tests and Holm–Sidak multiple comparisons tests (*p<0.05; **p<0.01; ***p<0.001; ****p<0.0001) All data are expressed as mean ± SD. A ROUT outlier test was performed and one data point corresponding to a Nrf2^+/+^ TXNRD1 GSR KO tumor in Supplementary Figure 3E–F for grade 1 burden was excluded.

## Acknowledgements

We would like to thank Dr. Edward Schmidt for providing the GSR KO and TXNRD1 flox mice. This work is supported by a grant from the Florida Department of Health Bankhead-Coley research program (9BC07) to G.M.D. This work has also been supported by the Lung Cancer Center of Excellence and Tissue Core and Microscopy Core Facilities at the Moffitt Cancer Center, an NCI designated Comprehensive Cancer Center (P30-CA076292).

## Author Contributions

Conceptualization, A.M.S and G.M.D.; Methodology, A.M.S., and G.M.D.; Investigation, A.M.S., J.M.D., S.C.; Writing – Original Draft, A.M.S and G.M.D.; Writing – Review & Editing, J.M.D and S.C.; Funding Acquisition, G.M.D; Supervision, G.M.D.

## Materials & Correspondence

Correspondence and material requests should be addressed to.

---

**Supplementary Figure 1.**
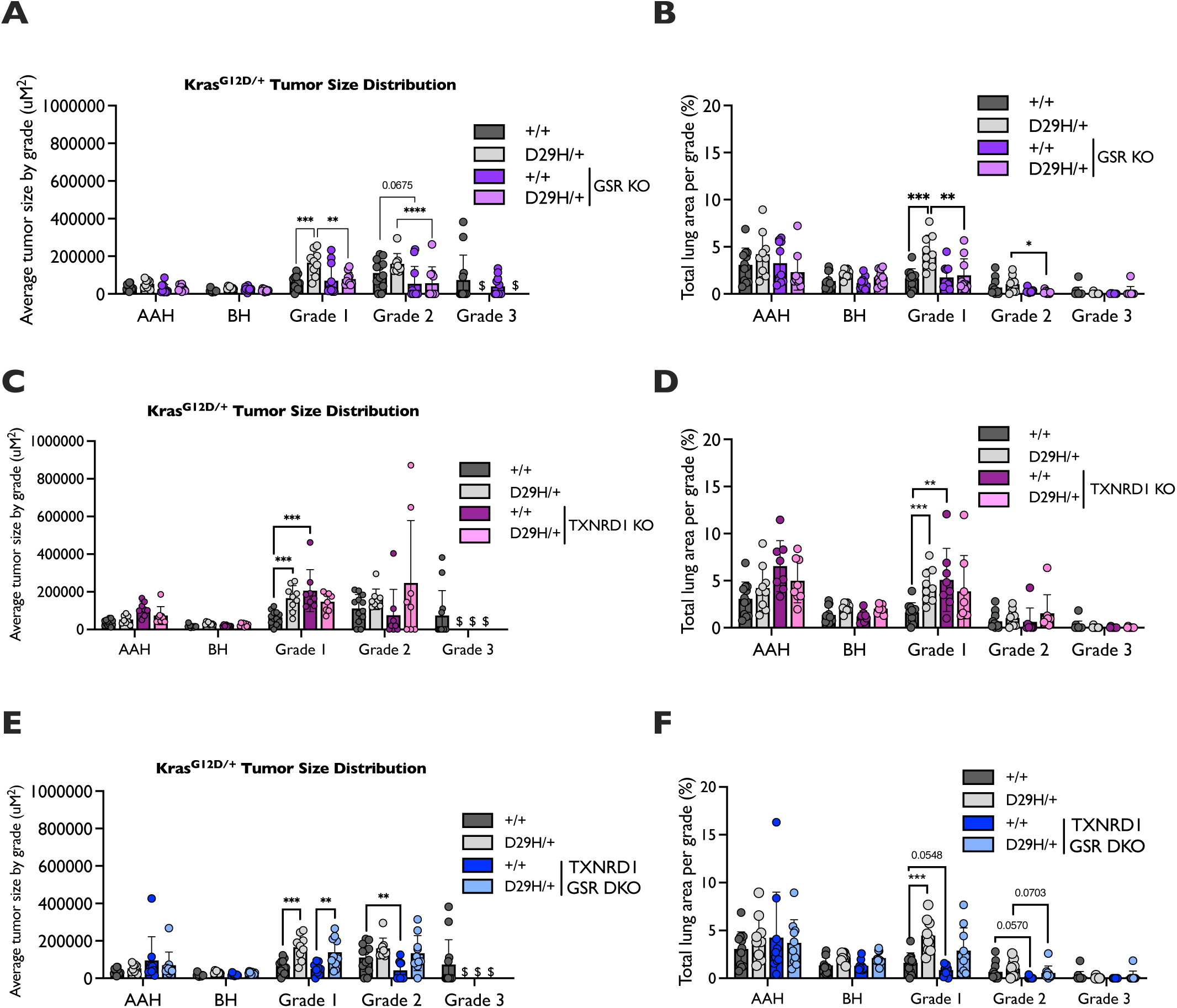
Lung tumor size analysis. Distribution of lung tumor size by grade across Kras^G12D/+^ and Kras^G12D/+;^ Nrf2^D29H/+^ mice for **(A)** GSR KO mice (*n* = 11 for Nrf2^+/+^; *n* = 10 for Nrf2^D29H/+^) **(C)** TXNRD1 KO mice (*n* = 8 for Nrf2^+/+^; *n* = 8 for Nrf2^D29H/+^), and **(E)** GSR/TXNRD1 KO mice (*n* = 10 for Nrf2^+/+^ and Nrf2^D29H/+^), compared to GSR/TXNRD1 WT mice (*n* = 12 for Nrf2^+/+^; *n* = 9 for Nrf2^D29H/+^, included in graphs in **A**, **C** and **E)**. Fraction of lung tumor burden by grade (lung tumor area per grade/total lung area) for **(B)** GSR KO mice (*n* = 11 for Nrf2^+/+^; *n* = 10 for Nrf2^D29H/+^), **(D)** TXNRD1 KO mice (*n* = 8 for Nrf2^+/+^; *n* = 8 for Nrf2^D29H/+^), and **(F)** GSR/TXNRD1 KO mice (*n* = 10 for Nrf2^+/+^ and Nrf2^D29H/+^), compared to GSR/TXNRD1 WT mice (*n* = 12 for Nrf2^+/+^; *n* = 9 for Nrf2^D29H/+^, included in graphs in **B**, **D** and **F**). *p<0.05, **p<0.01, ***p<0.001, ****p<0.0001 (unpaired t-test with Holm–Sidak multiple comparisons test). $ = fewer than three tumors detected across all mice. AAH, alveolar adenomatous hyperplasia; BH, bronchiolar hyperplasia.

**Supplementary Figure 2.**
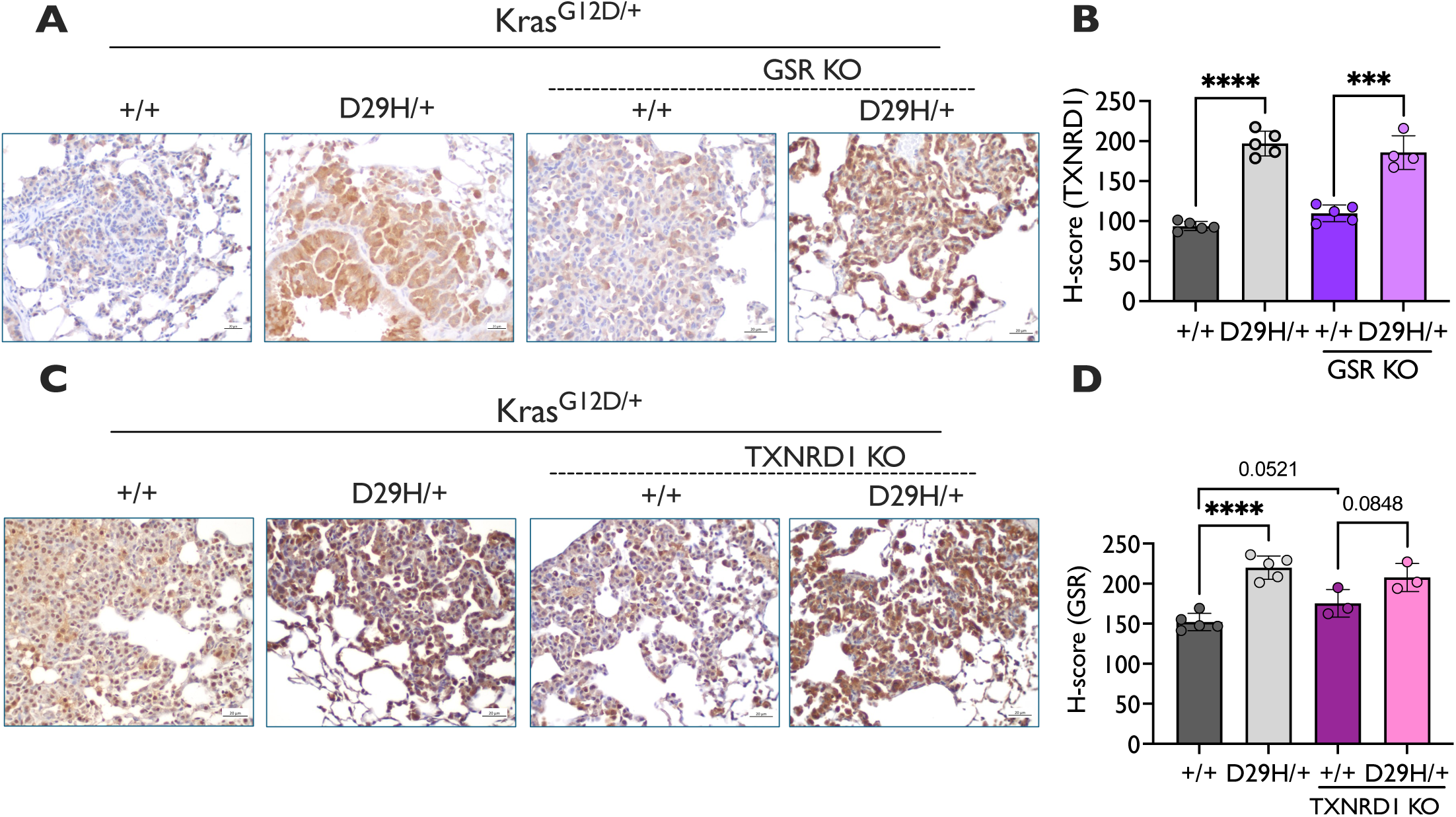
GSR or TXNRD1 KO does not increase GSR or TXNRD1 expression. **(A)** Representative TXNRD1 IHC staining of Nrf2 +/+ and D29H/+ tumors that are GSR WT or GSR KO and **(B)** quantitative H-score analysis for TXNRD1 staining. ****p<0.0001. Unpaired t-test. *n =* 5 for Nrf2^+/+^ and Nrf2^D29H/+^ GSR WT; Nrf2^+/+^ GSR KO (*n* = 5); Nrf2^D29H/+^ GSR KO (*n* = 4). Scale bars, 20 μm. **(C)** Representative GSR immunohistochemical (IHC) staining of Nrf2 +/+ and D29H/+ tumors that are TXNRD1 WT or TXNRD1 KO and **(D)** quantitative H-score analysis for GSR staining. ****p<0.0001. Unpaired t-test. *n =* 5 for Nrf2^+/+^ and Nrf2^D29H/+^ TXNRD1 WT; *n* = 3 for Nrf2^+/+^ and Nrf2^D29H/+^ TXNRD1 KO. Scale bars, 20 μm.

**Supplementary Figure 3.**
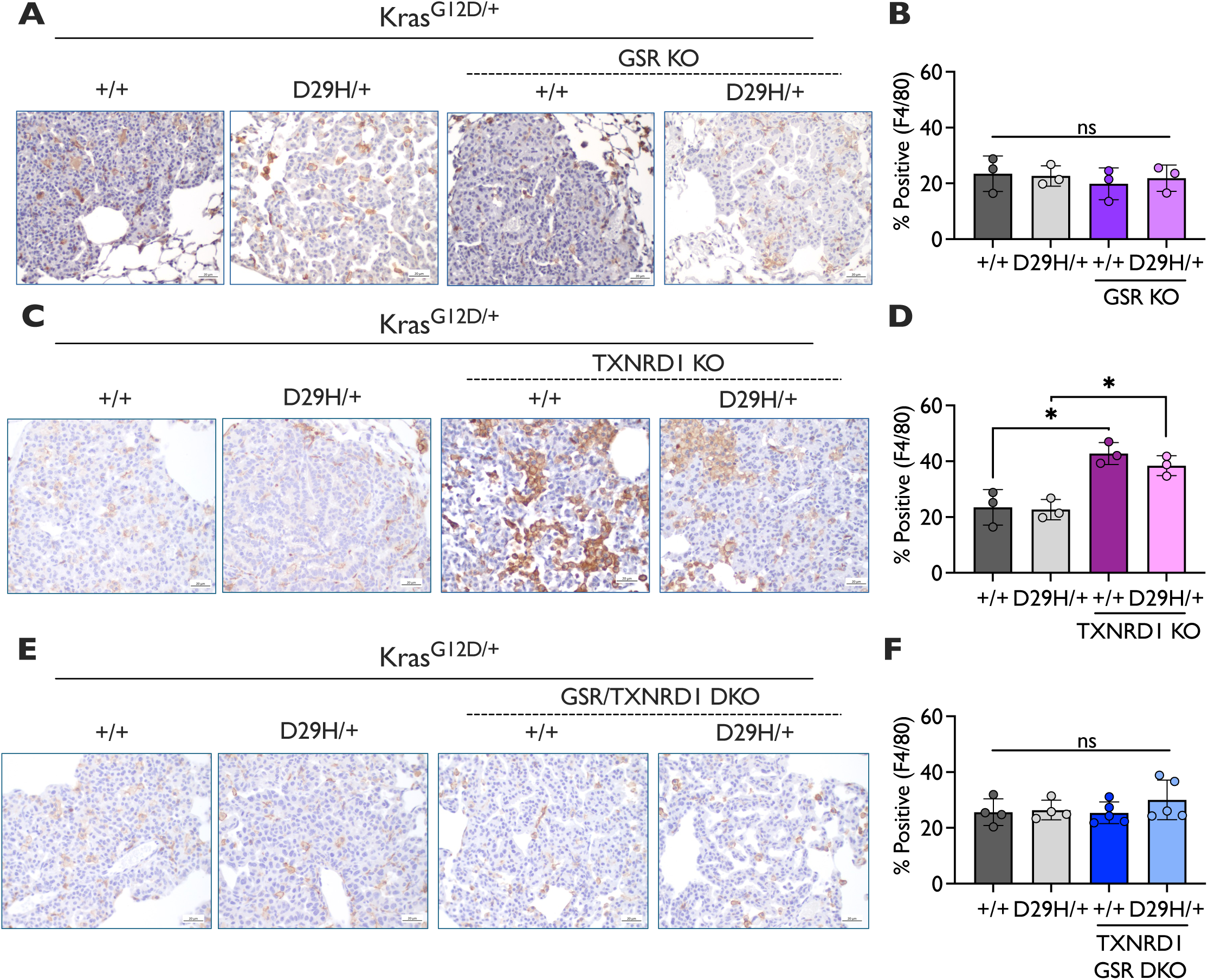
Macrophage infiltration is increased in TXNRD1 KO tumors. Representative IHC staining for the macrophage marker F4/80 of Nrf2^+/+^ and Nrf2^D29H/+^ tumors that are **(A)** GSR WT or GSR KO **(C)** TXNRD1 WT or TXNRD1 KO and **(E)** GSR/TXNRD1 WT or GSR/TXNRD1 KO. Percentage of F4/80 positive cells per tumor for (**B)** GSR KO mice (*n* = 3 for Nrf2^+/+^ and Nrf2^D29H/+^), **(D)** TXNRD1 KO mice (*n* = 3 for Nrf2^+/+^ and Nrf2^D29H/+^), and **(F)** GSR/TXNRD1 KO mice (*n* = 5 for Nrf2^+/+^ and Nrf2^D29H/+^), compared to GSR/TXNRD1 WT mice (*n* = 3 for Nrf2^+/+^ and Nrf2^D29H/+^, included in graphs in **B** and **D;** *n* = 4 for Nrf2^+/+^ and Nrf2^D29H/+^ in graph **F**). *p<0.05. Unpaired t-test).

## References

1 DeBlasi, J. M. & DeNicola, G. M. Dissecting the Crosstalk between NRF2 Signaling and Metabolic Processes in Cancer. Cancers (Basel) 12 (2020). 10.3390/cancers12103023

2 Kensler, T. W., Wakabayash, N. & Biswal, S. Cell survival responses to environmental stresses via the Keap1-Nrf2-ARE pathway. Annu Rev Pharmacol 47, 89–116 (2007). 10.1146/annurev.pharmtox.46.120604.141046

3 He, F., Ru, X. & Wen, T. NRF2, a Transcription Factor for Stress Response and Beyond. Int J Mol Sci 21 (2020). 10.3390/ijms21134777

4 Itoh, K. et al. Keap1 represses nuclear activation of antioxidant responsive elements by Nrf2 through binding to the amino-terminal Neh2 domain. Genes Dev 13, 76–86 (1999). 10.1101/gad.13.1.76

5 Dayalan Naidu, S. & Dinkova-Kostova, A. T. KEAP1, a cysteine-based sensor and a drug target for the prevention and treatment of chronic disease. Open Biol 10, 200105 (2020). 10.1098/rsob.200105

6 Hirotsu, Y. et al. Nrf2-MafG heterodimers contribute globally to antioxidant and metabolic networks. Nucleic Acids Res 40, 10228–10239 (2012). 10.1093/nar/gks827

7 Hayes, J. D. & Dinkova-Kostova, A. T. The Nrf2 regulatory network provides an interface between redox and intermediary metabolism. Trends Biochem Sci 39, 199–218 (2014). 10.1016/j.tibs.2014.02.002

8 Wu, K. C., Cui, J. Y. & Klaassen, C. D. Effect of graded Nrf2 activation on phase-I and - II drug metabolizing enzymes and transporters in mouse liver. PLoS One 7, e39006 (2012). 10.1371/journal.pone.0039006

9 He, F., Antonucci, L. & Karin, M. NRF2 as a regulator of cell metabolism and inflammation in cancer. Carcinogenesis 41, 405–416 (2020). 10.1093/carcin/bgaa039

10 Mitsuishi, Y. et al. Nrf2 Redirects Glucose and Glutamine into Anabolic Pathways in Metabolic Reprogramming. Cancer Cell 22, 66–79 (2012). 10.1016/j.ccr.2012.05.016

11 Kang, Y. P. et al. Cysteine dioxygenase 1 is a metabolic liability for non-small cell lung cancer. Elife 8 (2019). 10.7554/eLife.45572

12 Wu, K. C., Cui, J. Y. & Klaassen, C. D. Beneficial Role of Nrf2 in Regulating NADPH Generation and Consumption. Toxicol Sci 123, 590–600 (2011). 10.1093/toxsci/kfr183

13 Ludtmann, M. H. R., Angelova, P. R., Zhang, Y., Abramov, A. Y. & Dinkova-Kostova, A. T. Nrf2 affects the efficiency of mitochondrial fatty acid oxidation. Biochem J 457, 415–424 (2014). 10.1042/BJ20130863

14 Romero, R. et al. Keap1 loss promotes Kras-driven lung cancer and results in dependence on glutaminolysis. Nat Med 23, 1362-+ (2017). 10.1038%2Fnm.4407

15 Reisman, S. A., Yeager, R. L., Yamamoto, M. & Klaassen, C. D. Increased Nrf2 Activation in Livers from Keap1-Knockdown Mice Increases Expression of Cytoprotective Genes that Detoxify Electrophiles more than those that Detoxify Reactive Oxygen Species. Toxicol Sci 108, 35–47 (2009). 10.1093/toxsci/kfn267

16 Harvey, C. J. et al. Nrf2-regulated glutathione recycling independent of biosynthesis is critical for cell survival during oxidative stress. Free Radical Bio Med 46, 443–453 (2009). 10.1016/j.freeradbiomed.2008.10.040

17 Erickson, A. M., Nevarea, Z., Gipp, J. J. & Mulcahy, R. T. Identification of a variant antioxidant response element in the promoter of the human glutamate-cysteine ligase modifier subunit gene - Revision of the ARE consensus sequence. J Biol Chem 277, 30730–30737 (2002). 10.1074/jbc.m205225200

18 Deponte, M. Glutathione catalysis and the reaction mechanisms of glutathione-dependent enzymes. Bba-Gen Subjects 1830, 3217–3266 (2013). 10.1016/j.bbagen.2012.09.018

19 Moinova, H. R. & Mulcahy, R. T. Up-regulation of the human γ-glutamylcysteine synthetase regulatory subunit gene involves binding of Nrf-2 to an electrophile responsive element. Biochem Bioph Res Co 261, 661–668 (1999). 10.1006/bbrc.1999.1109

20 Hawkes, H. J. K., Karlenius, T. C. & Tonissen, K. F. Regulation of the human thioredoxin gene promoter and its key substrates: A study of functional and putative regulatory elements. Bba-Gen Subjects 1840, 303–314 (2014). 10.1016/j.bbagen.2013.09.013

21 Sakurai, A. et al. Transcriptional regulation of thioredoxin reductase 1 expression by cadmium in vascular endothelial cells: Role of NF-E2-related factor-2. J Cell Physiol 203, 529–537 (2005). 10.1002/jcp.20246

22 Osburn, W. O. et al. Nrf2 regulates an adaptive response protecting against oxidative damage following diquat-mediated formation of superoxide anion. Arch Biochem Biophys 454, 7–15 (2006). 10.1016/j.abb.2006.08.005

23 Salazar, M., Rojo, A. I., Velasco, D., de Sagarra, R. M. & Cuadrado, A. Glycogen synthase kinase-3β inhibits the xenobiotic and antioxidant cell response by direct phosphorylation and nuclear exclusion of the transcription factor Nrf2. J Biol Chem 281, 14841–14851 (2006). 10.1074/jbc.m513737200

24 Toppo, S., Flohé, L., Ursini, F., Vanin, S. & Maiorino, M. Catalytic mechanisms and specificities of glutathione peroxidases: Variations of a basic scheme. Bba-Gen Subjects 1790, 1486–1500 (2009). 10.1016/j.bbagen.2009.04.007

25 Flohé, L., Toppo, S., Cozza, G. & Ursini, F. A Comparison of Thiol Peroxidase Mechanisms. Antioxid Redox Sign 15, 763–780 (2011). 10.1089/ars.2010.3397

26 Rhee, S. G., Woo, H. A., Kil, I. S. & Bae, S. H. Peroxiredoxin Functions as a Peroxidase and a Regulator and Sensor of Local Peroxides. J Biol Chem 287, 4403–4410 (2012). 10.1074/jbc.r111.283432

27 Lu, J. & Holmgren, A. The thioredoxin antioxidant system. Free Radical Bio Med 66, 75–87 (2014). 10.1016/j.freeradbiomed.2013.07.036

28 Sun, Q. A. et al. Redox regulation of cell signaling by selenocysteine in mammalian thioredoxin reductases. J Biol Chem 274, 24522–24530 (1999). 10.1074/jbc.274.35.24522

29 Holmgren, A. Antioxidant Function of Thioredoxin and Glutaredoxin Systems. Antioxid Redox Sign 2, 811–U209 (2000). 10.1089/ars.2000.2.4-811

30 Holmgren, A. & Bjornstedt, M. Thioredoxin and Thioredoxin Reductase. Method Enzymol 252, 199–208 (1995). 10.1016/0076-6879(95)52023-6

31 Bannai, S. Exchange of Cystine and Glutamate across Plasma-Membrane of Human-Fibroblasts. J Biol Chem 261, 2256–2263 (1986).

32 Mandal, P. K. et al. System xc-and Thioredoxin Reductase 1 Cooperatively Rescue Glutathione Deficiency. J Biol Chem 285, 22244–22253 (2010). 10.1074%2Fjbc.M110.121327

33 Espinosa, B. & Arnér, E. S. J. Thioredoxin-related protein of 14 kDa as a modulator of redox signalling pathways. Brit J Pharmacol 176, 544–553 (2019). 10.1111/bph.14479

34 Pader, I. et al. Thioredoxin-related protein of 14 kDa is an efficient L-cystine reductase and S-denitrosylase. P Natl Acad Sci USA 111, 6964–6969 (2014). 10.1073/pnas.1317320111

35 Jiang, X. J., Stockwell, B. R. & Conrad, M. Ferroptosis: mechanisms, biology and role in disease. Nat Rev Mol Cell Bio 22, 266–282 (2021). 10.1038%2Fs41580-020-00324-8

36 Ingold, I. et al. Selenium Utilization by GPX4 Is Required to Prevent Hydroperoxide-Induced Ferroptosis. Cell 172, 409-+ (2018). 10.1016/j.cell.2017.11.048

37 Veal, E. A., Day, A. M. & Morgan, B. A. Hydrogen peroxide sensing and signaling. Mol Cell 26, 1–14 (2007). 10.1016/j.molcel.2007.03.016

38 Kamata, H. & Hirata, H. Redox regulation of cellular signalling. Cell Signal 11, 1–14 (1999). 10.1016/S0898-6568(98)00037-0

39 Zhang, Y. et al. Redox regulation of tumor suppressor PTEN in cell signaling. Redox Biol 34 (2020). 10.1016%2Fj.redox.2020.101553

40 Ghezzi, P. Regulation of protein function by glutathionylation. Free Radical Res 39, 573–580 (2005). 10.1080/10715760500072172

41 Best, S. A. et al. Synergy between the KEAP1/NRF2 and PI3K Pathways Drives Non-Small-Cell Lung Cancer with an Altered Immune Microenvironment. Cell Metab 27, 935-+ (2018). 10.1016/j.cmet.2018.02.006

42 Best, S. A. et al. Distinct initiating events underpin the immune and metabolic heterogeneity of *Kras*-mutant lung adenocarcinoma. Nat Commun 10 (2019). 10.1038/s41467-019-12164-y

43 Hayashi, M. et al. Microenvironmental Activation of Nrf2 Restricts the Progression of Nrf2-Activated Malignant Tumors. Cancer Res 80, 3331–3344 (2020). 10.1158/0008-5472.can-19-2888

44 Jeong, Y. et al. Role of KEAP1/NRF2 and TP53 mutations in lung squamous cell carcinoma development and radiation resistance. Cancer Res 77 (2017). 10.1158/2159-8290.cd-16-0127

45 Singh, A. et al. NRF2 Activation Promotes Aggressive Lung Cancer and Associates with Poor Clinical Outcomes. Clin Cancer Res 27, 877–888 (2021). 10.1158/1078-0432.CCR-20-1985

46 Lignitto, L. et al. Nrf2 Activation Promotes Lung Cancer Metastasis by Inhibiting the Degradation of Bach1. Cell 178, 316-+ (2019). 10.1016/j.cell.2019.06.003

47 Foggetti, G. et al. Genetic Determinants of EGFR-Driven Lung Cancer Growth and Therapeutic Response. Cancer Discov 11, 1736–1753 (2021). 10.1158/2159-8290.cd-20-1385

48 Rogers, Z. N. et al. Mapping the in vivo fitness landscape of lung adenocarcinoma tumor suppression in mice. Nat Genet 50, 483-+ (2018). 10.1038/s41588-018-0083-2

49 DeBlasi, J. M. et al. Distinct Nrf2 Signaling Thresholds Mediate Lung Tumor Initiation and Progression. Cancer Res 83, 1953–1967 (2023). 10.1158/0008-5472.can-22-3848

50 Jackson, E. L. et al. Analysis of lung tumor initiation and progression using conditional expression of oncogenic *K-ras*. Gene Dev 15, 3243–3248 (2001). 10.1101/gad.943001

51 Rogers, L. K. et al. Analyses of glutathione reductase hypomorphic mice indicate a genetic knockout. Toxicol Sci 82, 367–373 (2004). 10.1093/toxsci/kfh268

52 Pretsch, W. Glutathione reductase activity deficiency in homozygous mice does not cause haemolytic anaemia. Genet Res 73, 1–5 (1999). 10.1017/s0016672398003590

53 Bondareva, A. A. et al. Effects of thioredoxin reductase-1 deletion on embryogenesis and transcriptome. Free Radical Bio Med 43, 911–923 (2007). 10.1016%2Fj.freeradbiomed.2007.05.026

54 Harris, I. S. et al. Glutathione and Thioredoxin Antioxidant Pathways Synergize to Drive Cancer Initiation and Progression (vol 27, pg 211, 2015). Cancer Cell 27, 314–314 (2015). 10.1016/j.ccell.2014.11.019

55 Smith, J. A. Regulation of Cytokine Production by the Unfolded Protein Response; Implications for Infection and Autoimmunity. Front Immunol 9 (2018). 10.3389/fimmu.2018.00422

56 Poet, G. J. et al. Cytosolic thioredoxin reductase 1 is required for correct disulfide formation in the ER. Embo J 36, 693–702 (2017). 10.15252/embj.201695336

57 Conroy, L. R., Hawkinson, T. R., Young, L. E. A., Gentry, M. S. & Sun, R. C. Emerging roles of N-linked glycosylation in brain physiology and disorders. Trends Endocrinol Metab 32, 980–993 (2021). 10.1016/j.tem.2021.09.006

58 Nishinaka, Y., Nakamura, H. & Yodoi, J. Thioredoxin cytokine action. Protein Sensors and Reactive Oxygen Species, Pt a, Selenoproteins and Thioredoxin 347, 332–338 (2002). 10.1016/s0076-6879(02)47033-4

59 Rubartelli, A., Bajetto, A., Allavena, G., Wollman, E. & Sitia, R. Secretion of Thioredoxin by Normal and Neoplastic-Cells through a Leaderless Secretory Pathway. J Biol Chem 267, 24161–24164 (1992). 10.1016/S0021-9258(18)35742-9

60 Bertini, R. et al. Thioredoxin, a redox enzyme released in infection and inflammation, is a unique chemoattractant for neutrophils, monocytes, and T cells. J Exp Med 189, 1783–1789 (1999). 10.1084/jem.189.11.1783

61 Eriksson, S., Prigge, J. R., Talago, E. A., Arner, E. S. & Schmidt, E. E. Dietary methionine can sustain cytosolic redox homeostasis in the mouse liver. Nat Commun 6, 6479 (2015). 10.1038/ncomms7479

62 Yoon, S. J. et al. Comprehensive Metabolic Tracing Reveals the Origin and Catabolism of Cysteine in Mammalian Tissues and Tumors. Cancer Res 83, 1426–1442 (2023). 10.1158/0008-5472.CAN-22-3000

63 Weiss-Sadan, T. et al. NRF2 activation induces NADH-reductive stress, providing a metabolic vulnerability in lung cancer. Cell Metab 35, 487-+ (2023). 10.1016/j.cmet.2023.01.012

64 Sayin, V. I. et al. Activation of the NRF2 antioxidant program generates an imbalance in central carbon metabolism in cancer. Elife 6 (2017). 10.7554/eLife.28083

65 Muzumdar, M. D., Tasic, B., Miyamichi, K., Li, L. & Luo, L. Q. A global double-fluorescent cre reporter mouse. Genesis 45, 593–605 (2007). 10.1002/dvg.20335

